# Novel class of OTU deubiquitinases regulate substrate ubiquitination upon *Legionella* infection

**DOI:** 10.1101/2020.04.25.060954

**Authors:** Donghyuk Shin, Anshu Bhattacharya, Yi-Lin Cheng, Marta Campos Alonso, Ahmad Reza Mehdipour, Gerbrand J. van der Heden van Noort, Huib Ovaa, Gerhard Hummer, Ivan Dikic

## Abstract

*Legionella pneumophila* is a gram-negative pathogenic bacterium that causes Legionaries’ disease. The *Legionella* genome codes more than 300 effector proteins able to modulate host-pathogen interactions during infection. Among them are also enzymes altering the host-ubiquitination system including bacterial ligases and deubiquitinases. In this study, based on homology-detection screening on 305 *Legionella* effector proteins, we identified two <u>***L***</u>*egionella* <u>**OT**</u>U-like deubiquitinases (**LOT**; LotB (Lpg1621/Ceg23) and LotC (Lpg2529), LotA (Lpg2248/Lem21) is already known). A crystal structure of LotC catalytic core (LotC14-310) was determined at 2.4 Å and compared with other OTU deubiquitinases, including LotB. Unlike the classical OTU-family, the structures of *Legionella* OTU-family (LotB and LotC) shows an extended helical lobe between the Cys-loop and the variable loop, which define a novel class of OTU-deubiquitinase. Despite structural differences in their helical lobes, both LotB and LotC interact with ubiquitin. LotB has an additional ubiquitin binding site (S1’) enabling specific cleavage of Lys63-linked poly-ubiquitin chains. By contrast, LotC only contains the S1 site and cleaves different species of ubiquitin chains. MS analysis of catalytically inactive LotB and LotC identified different categories of host-substrates for these two related DUBs. Together, our results provide new structural insights of bacterial OTU deubiquitinases and indicate distinct roles of bacterial deubiquitinases in host-pathogen interactions.

## INTRODUCTION

Ubiquitination, a well-studied post-translational modification system, regulates the fate of various substrates by tagging them with ubiquitin (Yau and Rape, 2016). A representative example of ubiquitin-mediated cellular process is the ubiquitin-proteasome system (UPS), where misfolded proteins get ubiquitinated and degraded by the proteasome, and recycled (Dikic, 2017). For the larger cellular wastes, such as cellular components (ER, mitochondria etc.), protein aggregates or intracellular bacteria, ubiquitination works together with the autophagy machinery, which includes sequestration of ubiquitinated components and their transfer into the lysosome for degradation (Pohl and Dikic, 2019). To maintain homeostasis in the cell, ubiquitination events are tightly regulated by a reverse process called deubiquitination, where ubiquitin molecules from target substrates are specifically cleaved and recycled by deubiquitinating enzymes (deubiquitinases; DUBs) (Clague et al., 2019).

To date, about 100 different deubiquitinases (DUBs) have been identified from human. They are categorized into seven different classes, based on their structure and mechanism of actions including USP, JAMM (MPN), OTU, MJD (Josepin), UCH and the recently discovered MINDY and ZUFSP (Abdul Rehman et al., 2016; Clague et al., 2019; Haahr et al., 2018; Hermanns et al., 2018; Hewings et al., 2018; Kwasna et al., 2018). Six of them belong to the cysteine protease family (USP, OTU, MJD(Josepin), UCH, MINDY and ZUFSP), while JAMM (MPN) belongs to the zinc-containing metalloproteases. Among them, the OTU-family is distinguished from other DUBs, as they exhibit linkage specificity (Mevissen et al., 2016, 2013; Mevissen and Komander, 2017). For example, Cezanne specifically cleaves Lys11-linked polyubiquitin chains (Bremm et al., 2010) and OTUB1 preferentially cleaves Lys48-linked chains (Edelmann et al., 2009; Wang et al., 2009), while OTULIN exclusively cleaves M1-linked (linear) chains (Keusekotten et al., 2013). Extensive biochemical and structural studies have provided general mechanisms of how the diverse linkage specificity can be achieved within the structurally similar OTU-family, pointing to the presence or absence of additional ubiquitin binding domains (UBDs), sequence variations on ubiquitinated substrates, or additional S1’/S2 binding sites (Mevissen et al., 2013).

Considering the importance of ubiquitin-mediated cellular pathways, it is not surprising that pathogens are armed with various weapons to hijack the host-ubiquitination system. For instance, *Salmonella typhimurium* encodes HECT type E3 ligase SopA (Diao et al., 2008; Fiskin et al., 2017; Lin et al., 2012) and *Legionella pneumophila* contains LubX and LegU1, which are similar to U-box and F-box containing E3 ligases, respectively (Ensminger and Isberg, 2010; Kubori et al., 2008; Quaile et al., 2015). In addition, bacterial pathogens also possess atypical ubiquitin ligases that do not belong to any of known E3 ligases, such as IpaH family (*Shigella*) or SidC/SdCA (*Legionella*) (Hsu et al., 2014; Wasilko et al., 2018). More recently, the SidE family (SdeA, SdeB, SdeC and SidC) of *Legionella* has been shown to mediate unconventional phosphoribosyl (PR) serine ubiquitination which is also tightly regulated by the meta-effector SidJ or PR-ubiquitin specific deubiquitinases (DupA and DupB) (Bhogaraju et al., 2019, 2016; Black et al., 2019; Kalayil et al., 2018; Qiu et al., 2016; Shin et al., 2020). Pathogenic bacteria encode not only ubiquitin ligases, but also deubiquitinases (Hermanns and Hofmann, 2019). The most studied bacterial deubiquitinases are CE-clan proteases, based on the MEROPS database classification, which cleave either ubiquitin or ubiquitin-like modifiers (SUMO1 or NEDD8) (Pruneda et al., 2016; Rawlings et al., 2018). In addition to CE-clan DUBs, bacteria and viruses also encode OTU-like deubiquitinases. Several structures from viral-OTUs revealed that they have a unique structure compared to those of known OTU family members (Akutsu et al., 2011; Capodagli et al., 2013; James et al., 2011; Lombardi et al., 2013; van Kasteren et al., 2013). OTUs from nairovirus Crimean-Congo hemorrhagic fever virus (CCHFV) and Dugbe virus (DUGV) have an extra β-hairpin in their S1 binding site. Papain-like protease (PLP2) domain of the artervirus equine arteritis virus (EAV) contains a zinc-finger that also interacts with ubiquitin. While viral OTUs have been studied, to date only three bacterial OTU-like DUBs have been identified. ChlaOTU from *Chlamydophila pneumoniae* contains an OTU-domain that cleaves both K48- and K63-linked polyubiquitin chains (Furtado et al., 2013). LotA (Lpg2248;Lem21), <u>***L***</u>*egionella* <u>**OT**</u>U-like deubiquitinase, contains two OTU-like domains with two catalytic Cys residues (C13, C303), where both are required for cleaving ubiquitin chains from an LCV (*Legionella*-containing vacuole) (Kubori et al., 2018). Interestingly, LotA showed K6-linkage preference that is solely dependent on its first OTU domain (Cys13). Recently, another OTU-like deubiquitinase from *Legionella* (Lpg1621, Ceg23) has been identified as K63 chain-specific OTU-deubiquitinase (Ma et al., 2020).

Despite these findings, little is known about the structure and molecular details of bacterial OTU-like deubiquitinases. Here, we describe the two novel OTU-like deubiquitinaseses in *Legionella* (LotB (Lpg1621; Ceg23) and LotC (Lpg2529). Structural analysis on LOT-deubiqutinases provides insights on how bacterial OTU-deubiquitnases distinguished from the known OTU members. Furthermore, we also identified the specific host-substrates of LotB and LotC by mass-spectrometry (MS) analysis using catalytically inactive variants. Collectively, our findings provide valuable structural insights into bacterial deubiquitinases and their roles in host-pathogen interactions.

## RESULTS

### Identification of two novel OTU-like deubiquitinases from *Legionella* effector proteins

To identify putative deubiquitinases amongst *Legionella* effector proteins, we analyzed effector proteins from *L. pneumophlia* (Lpg genes). Based on the type IV Icm/Dot complex secretion signal (> 2.0), 305 effector proteins were selected (Burstein et al., 2016). Using pairwise sequence-structure comparison based on hidden Markov models (HMMs, HHpred suite) (Zimmermann et al., 2018), revealed four previously uncharacterized proteins as putative deubiquitinases. These proteins all contain catalytic domains of known deubiquitinases (Fig. 1a). Lpg1621(Ceg23) and Lpg2529 are found as members of OTU-family, whereas Lpg2411 and Lpg2907 belong to the UCH and CE-clan, respectively (Fig. 1b, Table 1). An *in vitro* di-ubiquitin cleavage assay with di-Ub panel (eight different linkage-specific Ub2 chains(El Oualid et al., 2010)) showed that the OTU-like DUBs (Lpg1621 and Lpg2529) are capable of cleaving Ub2 chains with different specificity, while other candidates (Lpg2411 and Lpg2907) did not show catalytic activity (Supplementary Fig. 1a). The OTU family deubiquitinases have been shown to have linkage specificity against certain polyubiquitin chains (Mevissen et al., 2016, 2013). To address whether Lpg1621 and Lpg2529 follow this fundamental rule, we performed a time-course *in vitro* DUB assay with di-Ub panel (Fig. 1c-f). Consistent with the recent evidence, Lpg1621 exclusively processed the K63-linked Ub2 (Ma et al., 2020), while Lpg2529 showed activity against K6-, K11-, K33-, K48- and K63- linked Ub2. Based on the sequence homology and catalytic activity, we now re-named the Lpg1621 and Lpg2529 as <u>***L***</u>*egionella* <u>**OT**</u>U-like deubiquitinases (LotB and LotC, respectively).

**Figure 1.**
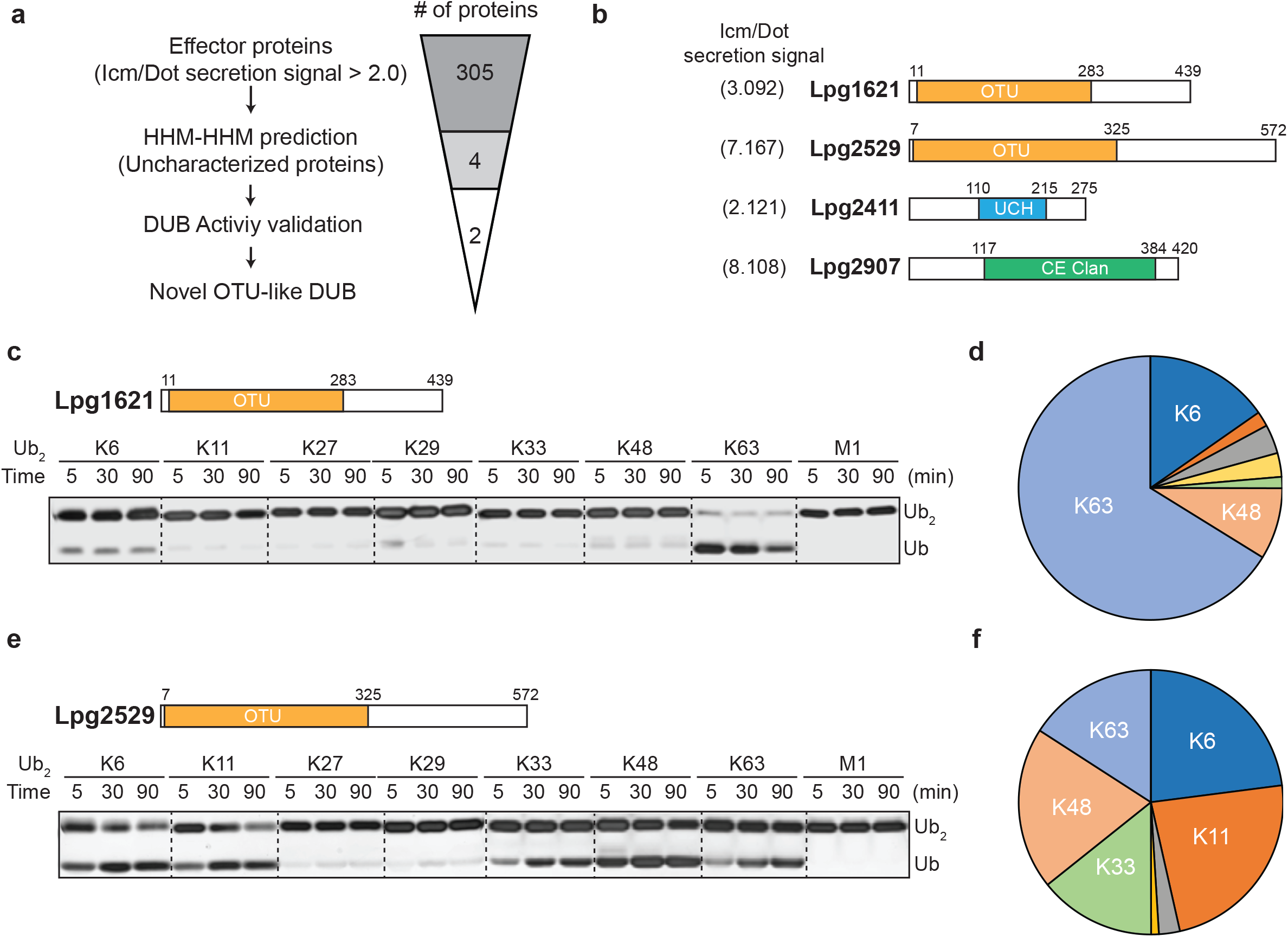
Identification of novel deubiqutinases in *Legionella pneumophila.* **(a)** Graphical illustration of identification of novel deubiquitinases of Legionella pneumophila Lpg Genes. **(b)** Predicted deubiquitinase domain of four putative *Legionella* deubiquitinases. **(c, e)** Time-course Di-ubiquitin panel cleavage assay with Lpg1621 (LotB) and Lpg2529 (LotC), respectively. **(d, f)** Linkage specificity diagram of Lpg1621 (LotB) and Lpg2529 (LotC). Percentage of cleaved ubiquitin species at 90 mins were plotted.

**Table1.**
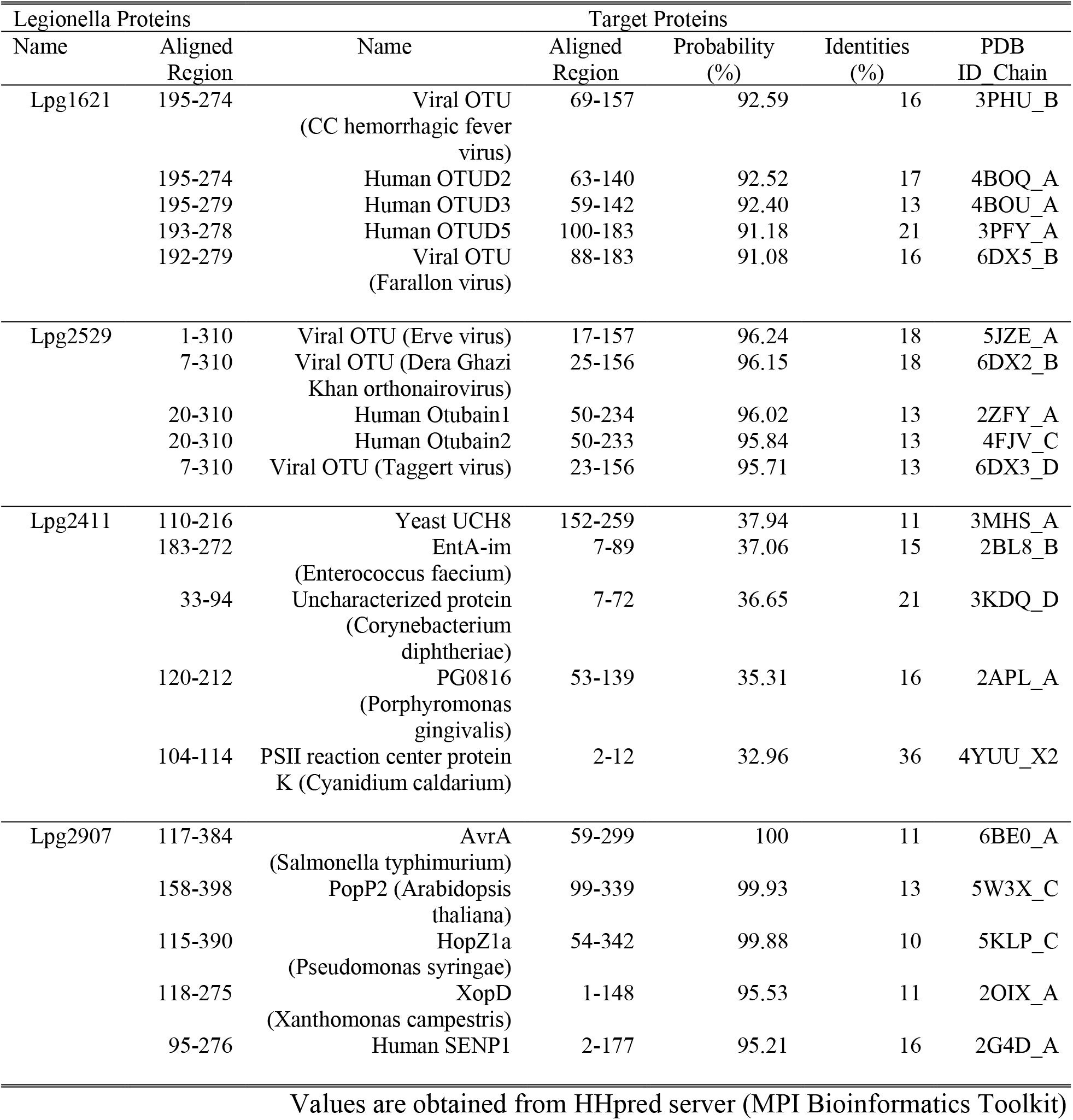
TOP 5 candidates for putative DUBs from Legionella effector proteins.

| Legionella Proteins |  | Target Proteins |  |  |  |  |
| --- | --- | --- | --- | --- | --- | --- |
| Name | Aligned Region | Name | Aligned Region | Probability (%) | Identities (%) | PDB ID Chain |
| Lpg1621 | 195-274 | Viral OTU<br>(CC hemorrhagic fever virus) | 69-157 | 92.59 | 16 | 3PHU_B |
|  | 195-274 | Human OTUD2 | 63-140 | 92.52 | 17 | 4BOQ_A |
|  | 195-279 | Human OTUD3 | 59-142 | 92.40 | 13 | 4BOU_A |
|  | 193-278 | Human OTUD5 | 100-183 | 91.18 | 21 | 3PFY_A |
|  | 192-279 | Viral OTU<br>(Farallon virus) | 88-183 | 91.08 | 16 | 6DX5_B |
| Lpg2529 | 1-310 | Viral OTU (Erve virus) | 17-157 | 96.24 | 18 | 5JZE_A |
|  | 7-310 | Viral OTU (Dera Ghazi Khan orthonairovirus) | 25-156 | 96.15 | 18 | 6DX2_B |
|  | 20-310 | Human Otubain1 | 50-234 | 96.02 | 13 | 2ZFY_A |
|  | 20-310 | Human Otubain2 | 50-233 | 95.84 | 13 | 4FJV_C |
|  | 7-310 | Viral OTU (Taggert virus) | 23-156 | 95.71 | 13 | 6DX3_D |
| Lpg2411 | 110-216 | Yeast UCH8 | 152-259 | 37.94 | 11 | 3MHS_A |
|  | 183-272 | EntA-im<br>(Enterococcus faecium) | 7-89 | 37.06 | 15 | 2BL8_B |
|  | 33-94 | Uncharacterized protein<br>(Corynebacterium diphtheriae) | 7-72 | 36.65 | 21 | 3KDQ_D |
|  | 120-212 | PG0816<br>(Porphyromonas gingivalis) | 53-139 | 35.31 | 16 | 2APL_A |
|  | 104-114 | PSII reaction center protein K (Cyanidium caldarium) | 2-12 | 32.96 | 36 | 4YUU_X2 |
| Lpg2907 | 117-384 | AvrA<br>(Salmonella typhimurium) | 59-299 | 100 | 11 | 6BE0_A |
|  | 158-398 | PopP2 (Arabidopsis thaliana) | 99-339 | 99.93 | 13 | 5W3X_C |
|  | 115-390 | HopZ1a<br>(Pseudomonas syringae) | 54-342 | 99.88 | 10 | 5KLP_C |
|  | 118-275 | XopD<br>(Xanthomonas campestris) | 1-148 | 95.53 | 11 | 2OIX_A |
|  | 95-276 | Human SENP1 | 2-177 | 95.21 | 16 | 2G4D_A |

Values are obtained from HHpred server (MPI Bioinformatics Toolkit)

### Biochemical properties of LotB and LotC

The OTU-family belongs to the cysteine protease family, requiring the presence of a catalytic triad for their activity (Mevissen et al., 2013). Based on the sequence analysis, we identifed the conserved catalytic triad for both LotB (D27, C29 and H270) and LotC (D17, C24 and H304). Mutations on either cysteine or histidine completely abolished the catalytic activity of both DUBs, which suggests that both LotB and LotC follow the general catalytic mechanism of the OTU-family (Fig. 2a, c). Next, we sought to find whether LotB and LotC require additional ubiquitin binding sites (S1’ or S2). To elicit this information, we used two different types of ubiquitin activity-based probes (ABP). The propargyl-ubiquitin-ABP (Prg-ABP) contains a highly reactive propargyl group at the C-terminus of ubiquitin chains, which can form a covalent bond with the catalytic cysteine (Ekkebus et al., 2013; Flierman et al., 2016; Sommer et al., 2013). The vinylmethylester-ubiquitin-ABP (VME-ABP) contains VME, which replaces the isopeptide bond between two ubiquitin moieties in chains, and also forms a covalent bond with the catalytic cysteine (Borodovsky et al., 2002; Mulder et al., 2014). Both LotB and LotC showed clear shifts with all Prg-ABPs (mono-, K48- and K63- linked) with different reactivity. LotB shifted after 30 mins, and only partially, as evident by the amount of unreacted species, while LotC has rapidly reacted with Prg-ABP and was completely conjugated after 30 minutes (Fig. 2d, e). These results suggest that both LotB and LotC have a primary ubiquitin binding S1 site where the propargyl group can be located in close proximity to the catalytic cysteine. In contrast with Prg-ABP, only LotB reacted with K63-Ub2-VME-ABP. This is consistent with the di-Ub panel assay (Fig. 1c-f), in which LotB showed specificity toward the K63-linkage. The VME-ABp results suggest an extra S1’ ubiquitin binding site on LotB, which helps to properly locate the K63-linked-VME group on the catalytic site proximal to the catalytic cysteine. LotC lacks this S1’ site so that VME group between two ubiquitin moieties cannot be reached to the catalytic cysteine. Next, we asked whether both LotB and LotC can also interact with other ubiquitin-like modifiers (SUMO1/2/3, NEDD8) (Fig. 2f, g, Supplementary Fig. 2a, b). Interestingly, LotB showed modification with both NEDD8-Prg and ubiquitin-Prg after 30 minutes of reaction, while LotC was modified only with ubiquitin. It suggests that LotB binds to ubiquitin through the conserved binding interface with NEDD8, while LotC interacts with ubiquitin through the specific residues present only in ubiquitin (Supplementary Fig. 2c).

**Figure 2.**
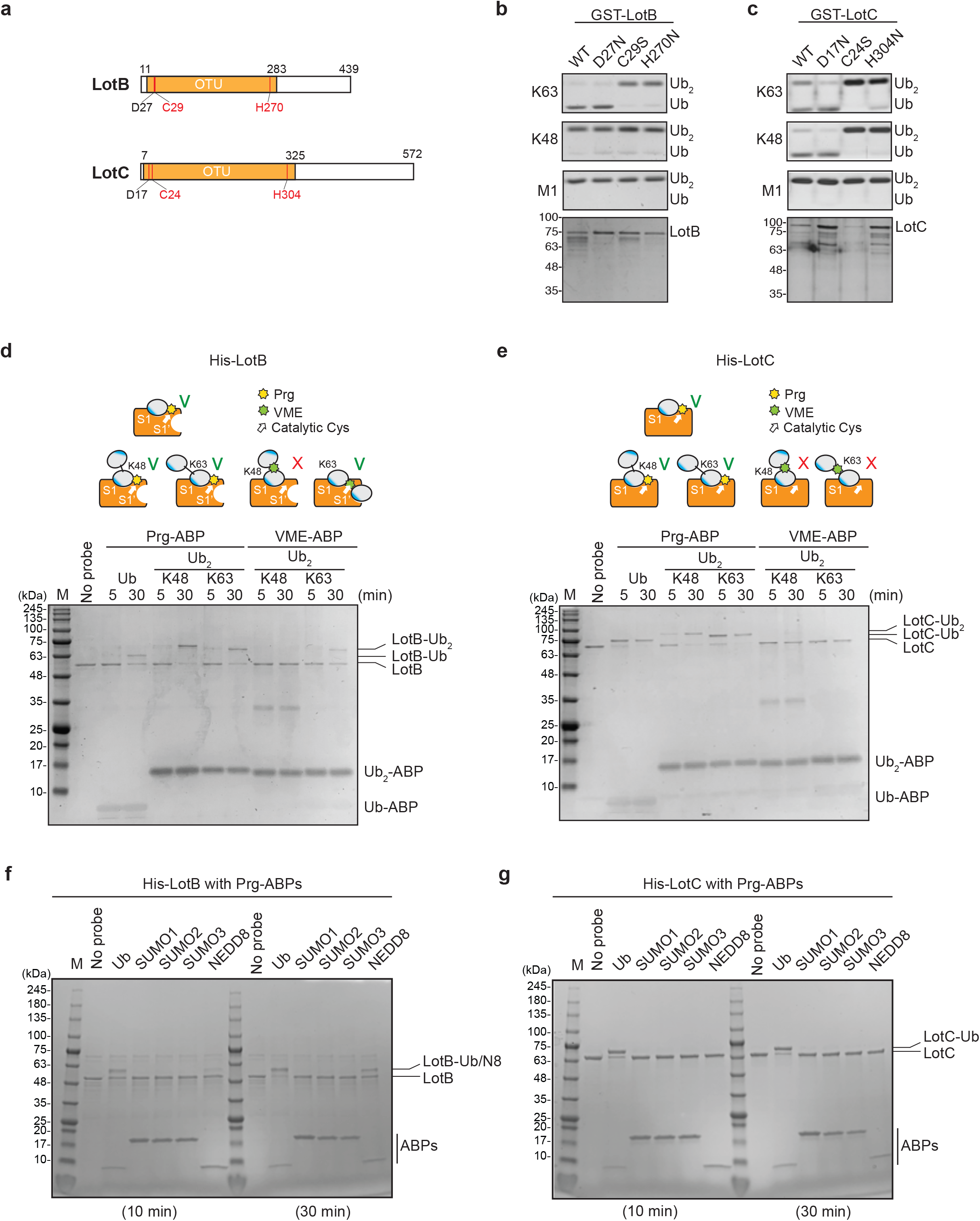
Biochemical properties of LotB and LotC. **(a)** Predicted catalytic residues on LotB and LotC. **(b, c)** Di-Ub cleavage activity assay with wild-type and catalytic mutants of LotB and LotC, respectively. **(d, e)** Activity based probes (ABPs) test on LotB and LotC. Propargyl-Ub-ABP (Prg-ABP) and vinylmethylester-ubiquitin-ABP (VME-ABP) were incubated as indicated time-points with LotB and LotC and analyzed on SDS-PAGE with Coomassie staining. **(f, g)** Propargyl ubiquitin or ubiquitin-like modifiers reactivity test on LotB and LotC. Prg-ABPs are incubated with LotB and LotC with indicated time points.

### Structural analysis of *Legionella* OTU-like deubiquitinases

Linkage specificity of the OTU family relies on one of the following mechanisms: 1) additional ubiquitin-binding domains, 2) ubiquitinated sequences in the substrates or 3) defined S1’ or S2 ubiquitin-binding sites (Mevissen et al., 2013). To define the minimal OTU domain for biochemical and structural studies, we designed several constructs and tested their activity against the di-Ub panel (Fig. 3a, b). While LotC retained its activity with the predicted OTU domain (7-310), LotB lost its activity after deletion of 50 amino acids (300-350) located at the C-terminus beyond the predicted OTU domain (11-283). Based on the LotB structure (PDB:6KS5, (Ma et al., 2020)), we assumed that this extra helical region might be required for the additional ubiquitin binding site (S1’) to accept the distal ubiquitin moiety from K63 Ub_2_ (Fig. 3c). To understand the detailed mechanism of linkage specificity of LotB and LotC at the molecular level, we determined the crystal structure of the catalytic domain of LotC (LotC_14-310_) at 2.4 Å (Fig. 3d, Supplementary Table 1, PDB ID: 6YK8). A structural comparison of both LotB and LotC with other OTU-deubiquitinases predicted by HHpred revealed that both Lot-DUBs have the unique structural features in the S1 ubiquitin-binding site (Fig. 3c, d and Table 1). Whereas the overall fold of the catalytic core of LotB and LotC resembles that of other OTU-deubiquitinases, both showed clear differences in the helical arm region, which has been shown to interact with ubiquitin and it serves as an S1 binding site (Mevissen et al., 2013). The structure and sequence alignment with other OTUs clearly showed that both LotB and LotC contain a relatively long insertion between the Cys-loop and the variable loop, compared to other OTU members (Fig. 3e). The typical length of the helical lobe of the known OTUs is ranging from 50 to 60 amino acids (except Otubain family which contain 110-120 amino acids), while LotB and LotC contain 183 and 210 amino acids, respectively. Based on this observation, we wondered whether LotA, another *Legionella* OTU-deubiquitinase (Kubori et al., 2018), also contains a longer insertion in the same region. Based on the catalytic cysteine and histidine residues of the two OTU domains on LotA (Hermanns and Hofmann, 2019), we analyzed the sequence and found that both OTU domains of LotA also contain the longer insertion between the Cys loop and the variable loop (179 and 178 amino acids, respectively; Fig. 3e). Together, our results identify Lot-DUBs as a novel class of the OTU-family with longer insertions in the helical lobe region (Supplementary Fig.3a).

**Figure 3.**
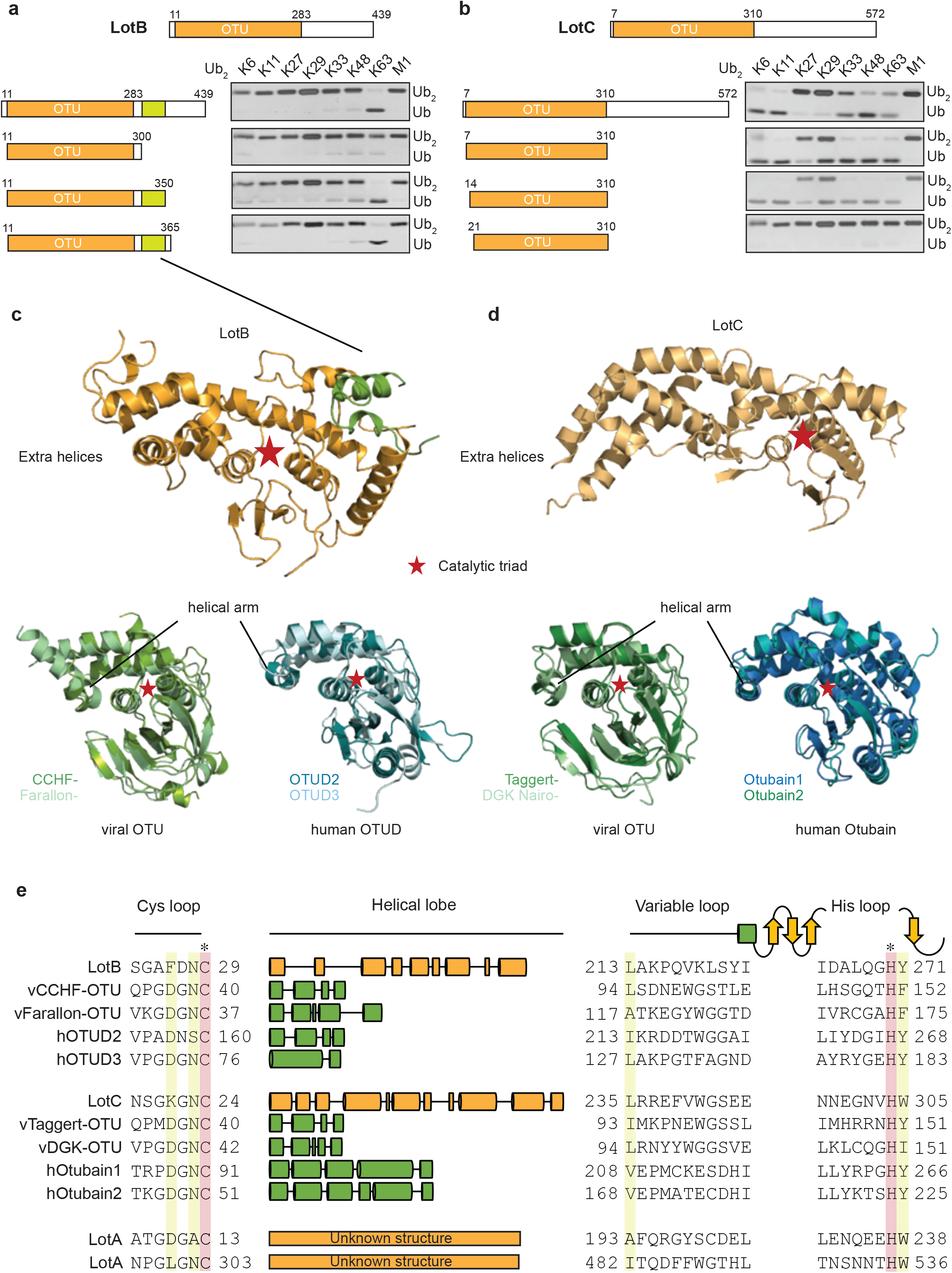
Structural comparison of *Legionella* OTU-DUBs with other OTU-family. **(a, b)** Minimal domain boundaries of catalytically active LotB and LotC. Different constructs were cloned based on the predicted OTU-domains and their activity, and were tested with di-Ub2 panel **(c, d)** Structural comparison of LotB and LotC with their closest homologues. CCHF- (PDB: 3PHU), Farallon- (PDB: 6DX5), OTUD2 (PDB: 4BOQ), OTUD3 (PDB: 4BOU), Taggert- (PDB: 6DX3), DGK nairo- (PDB: 6DX2), Otubain1 (PDB: 2ZFY), Otubain2 (PDB: 4FJV). **(e)** Sequence alignment of LotB and LotC with their closest homologues. Catalytic cysteine and histidine are highlighted in red and conserved residues are highlighted in yellow.

### Novel structural fold of S1-ubiquitin binding sites on *Legionella* OTUs

Both LotB and LotC have extended helices, specifically near the S1 ubiquitin binding site and we wondered how these regions interact with ubiquitin. To address this, we performed ubiquitin docking into both LotB and LotC, followed by molecular dynamics (MD) simulations for 600 ns (Fig. 4a-d). The final models showed that ubiquitin indeed makes contacts with the additional helical regions of both LotB and LotC. In LotB, Phe143 and Met144 make hydrophobic interactions with ubiquitin (Phe45 and Ala46). In addition to these interactions, we also found another hydrophobic patch in LotB (Ile238, Val247, Ala248, Ile264, Ala266) interacting with ubiquitin (Leu71, Leu73). For LotC, we identified several hydrophobic interactions of the extended helical region (Tyr119, Tyr149) with ubiquitin (Ile44). During the simulation, C-terminus of ubiquitin (Arg72, Arg74) formed transient electrostatic interactions with LotC (Glu153, Glu245). To validate the interactions observed in the simulations, we introduced several mutations to the binding interface of both LotB and LotC, and performed a ubiquitin-cleavage assay (Fig.4e, f). Consistent with a recent study(Ma et al., 2020), mutations of both F143 and M144 decreased the catalytic activity of LotB. Interestingly, mutations of the newly identified hydrophobic patch (I238, A266) also decreased the catalytic activity. For LotC, the mutations in the hydrophobic patch (Y119, Y149) also affected the catalytic activity of LotC. Remarkably, a single mutation on E153 completely abolished the catalytic activity, which indicates that the electrostatic interactions are important for the correct docking of the ubiquitin C-terminus to the catalytic pocket of LotC. This observation also explains the result observed in ubiquitin-like protein ABP assays with LotC (Fig. 2g). The NEDD8, which has alanine instead of Arg72 in ubiquitin, showed no modification towards LotC (Supplementary Fig. 2c). Together, our results reveal how the extra helical lobes of the Lot-DUBs interact with ubiquitin and how they differ within the *Legionella* OTU family.

**Figure 4.**
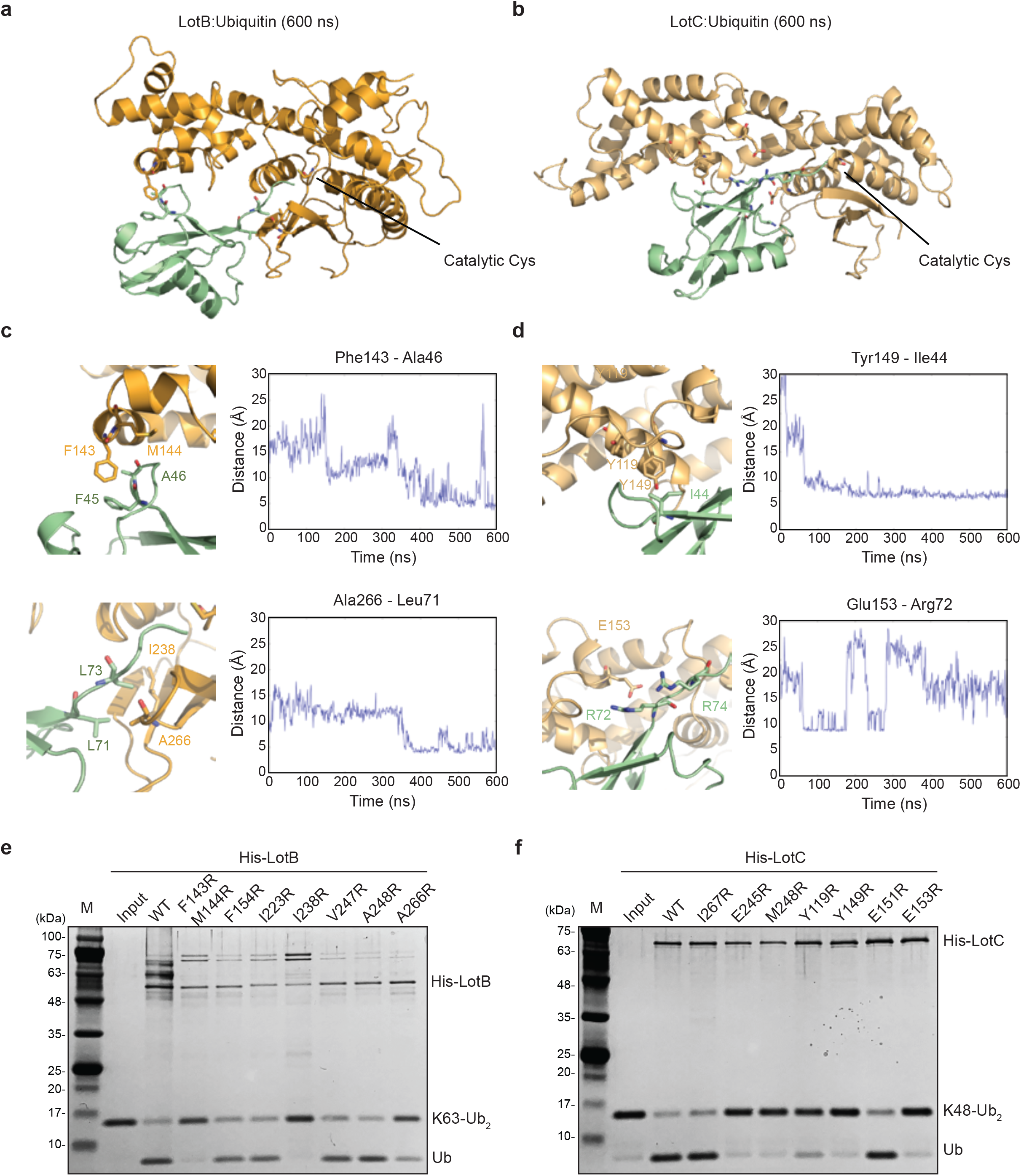
Ubiquitin binding sites on LotB and LotC. **(a, b)** Molecular docking and simulations of monoubiquitin to LotB and LotC, respectively. Shown are representative snapshots of the MD simulations. Catalytic cysteine and key residues for the interaction between ubiquitin and LotB or LorC are depicted as sticks. **(c, d)** Key residues mediating interactions of ubiquitin with LotB and LotC, respectively. Residues are highlighted in the structure (left). Side-chain center-of-mass distances are shown as a function of the simulation time (right). **(e, f)** Di-ubiquitin cleavage assay for mutants of LotB and LotC. The catalytic activity of LotB or LotC wild-type and its mutants was tested with K63- or K48- linked Ub2, respectively.

### Physiological roles of LotB and LotC

To gain better insights into the physiological roles of LotB and LotC, we decided to identify their interacting proteins or substrates. First, to enrich for the interacting partners, catalytically inactive LotB or LotC were expressed in cells and immuno-precipitated from cell lysates. Ubiquitin (UBA52) is strongly enriched with both catalytically inactive LotB and LotC (Fig. 5a, c). MS analysis revealed that LotB mainly interacts with membrane protein complexes (COPB1, ATP5B, ATP5H, COX5A, SEC61B). We also found interactions with some ER-resident proteins (Calnexin (CANX), DDOST, STT3A). By contrast, most of the enriched proteins from the inactive LotC pull-down were non-membrane-bound organelle-and ribosome-related proteins (RPS8, RPLP2, RPS27, RPLP1, RPL13) (Fig. 5a, c). To further understand this, we sought to find the cellular localization of both DUBs (Fig. 5b, d). Consistent with the recent publication, LotB specifically co-localized with the ER marker protein Calnexin, but not with other organelle markers (TOMM20 and GM130 for Mitochondria and Golgi, respectively, Fig. 5b), and the OTU domain itself failed to localize on the ER (Supplementary Fig. 4a). By contrast, we could not find a specific cellular localization of LotC (Fig. 5d). Next, to gain more insights into the functional roles of LotB and LotC, we decided to explore combinatorial ubiquitination events with other ubiquitin-related *Legionella* effector proteins. To do this, we co-transfected the cells with one of the Lot-DUBs and previously known *Legionella* E3 ligases (SidC or SdcA) (Hsu et al., 2014; Wasilko et al., 2018). We chose these two ligases because their cellular substrates are poorly studied. TMT-labelled samples from cells expressing either SidC or SdcA alone, or together with a catalytically inactive mutant of LotB or LotC, were prepared (four combinations; SidC-LotB, SidC-LotC, SdcA-LotC or SdcA-LotC, Fig. 5e, f, Supplementary Fig. 5a-d). Interestingly, we identified a distinct sub-class of substrates. Overall, a smaller number of proteins were enriched with LotB compared to LotC. We reasoned that LotB specifically interacts with proteins modified with K63-linked ubiquitin chains, while LotC interacts with different types of ubiquitin chains. Intriguingly, we found a significant number of ribosome-structural proteins in LotC:SdcA combination, which were not enriched in SidC background. Together, these results provide fundamental insights into how *Legionella*-derived E3-ligases and DUBs interact with each other.

**Figure 5.**
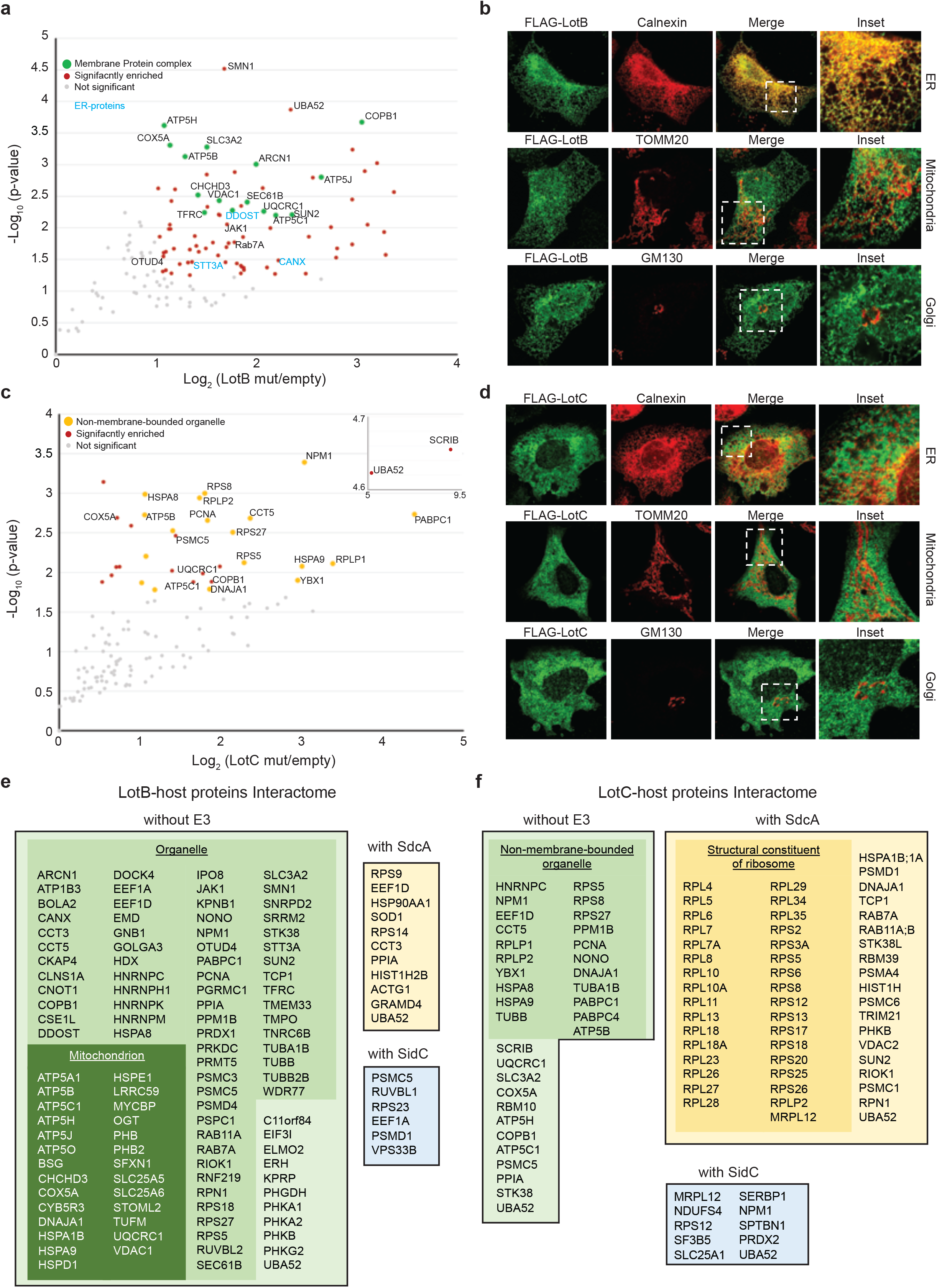
Physiological roles of LotB and LotC. **(a, c)** Proteomic analysis of interacting partners of LotB and LotC, respectively. Catalytically inactive FLAG-LotB (C29A) and FLAG-LotC (C24A) were transfected and immunoprecipitated. Co-precipitated interacting proteins were analyzed by mass spectrometry. **(b, d)** Cellular localization of LotB and LotC. FLAG-tagged LotB and LotC were ectopically expressed in U2OS cells and immunostained with cellular organelle markers (ER; Calnexin, Mitochondria: TOMM20, Golgi: GM130). **(e, f)** List of significantly enriched proteins from LotB and LotC with or without *Legionella* ligases (SdcA or SidC) as indicated.

## DISCUSSION

In this study, we have identified two novel bacterial OTU-deubiquitinases from *Legionella*, which we suggest as founding members of a new sub-class. Unlike classical OTU-DUBs, the *Legionella* OTU-DUBs (LOT) possess extended helical insertions between the catalytic Cys-loop and the variable loop. Molecular dynamic simulations with biochemical studies showed that the helical insertions interact with ubiquitin. As this insertion is unique for *Legionella* OTUs and not found in other known OTU-family members, *Legionella* OTU-DUBs define a new sub-class of the OTU-deubiquitinase family. We have also shown that LotB and LotC are specific to certain ubiquitin chains and have a distinct cellular localization. Moreover, host-protein interactome studies revealed that LotB and LotC have different sets of host-interacting proteins to LotB and LotC. Together, these findings establish important guidance on how to screen for more DUBs in other pathogenic bacteria or viruses, how to characterize their physiological roles during infection.

We also showed that the two *Legionella* OTUs have different ubiquitin-binding modes that enable them to cleave specific ubiquitin chains. With ubiquitin activity-based probes (Prg-, VME- probes), we showed that LotB contains an extra ubiquitin-binding site (S1’) and is specific to K63-linked ubiquitin chains, wherease LotC cleaves different types of ubiquitin chains. Interestingly, we observed a modification of LotB with NEDD8-Prg ABP. Further studies on neddylated proteins with LotB will give us more insights into dual-activity of LotB. In contrast, we could not see the modification between NEDD8-ABP and LotC and we reasoned that the Arg72 on ubiquitin, which is replaced by alanine in NEDD8, is important to locate the C-terminus of ubiquitin to the catalytic site. Indeed, in molecular dynamic simulations of LotC with ubiquitin we found Arg72 to from transient interactions with Glu153 of LotC. Further structural analysis will shed light on how different ubiquitin chains bind to LotB or LotC through the different binding modes should give us further insights.

Deubiquitinases from bacteria and viruses have been shown to alter the host immune system. For example, papain-like proteases (PLPro) from coronaviruses, such as middle east respiratory syndrome (MERS) or severe acute respiratory syndrome (SARS) have dual deubiquitinase and de-ISGylation activities and antagonized type I interferon (IFN-I) response, which is the primary defense system against viral infections (Davis and Gack, 2015; Devaraj et al., 2007; Frieman et al., 2009; Sadler and Williams, 2008). Interestingly, we identified OTUD4 as an interacting partner of LotB. OTUD4 has been shown to deubiquitinate K63-linked chains of myeoloid differentiation primary response 88 (MYD88) and to downregulate NF-kB-dependent inflammation. While the recent study on LotB showed that there is no detectable inhibition of NF-kB reporter expression(Ma et al., 2020), further studies are awaited to show the cross-talk between LotB and host ubiquitination system.

*Legionella pneumophila* has been shown to possess multiple genes altering the host ubiquitination system. However, little is known about functional cross-talk between ubiquitin ligases and deubiquitinases. To understand the combinatorial effects of *Legionella* ubiquitin ligases and deubiquitinases, we analyzed the interactome of LotB/LotC with host proteins in the presence or absence of *Legionella* ligases (SidC and SdcA). We could see clear differences in the number of enriched proteins in different combinations. Interestingly, we saw the clear difference in the number of enriched proteins. A significant number of ribosomal proteins were enriched with LotC in the background of SdcA, but not from SidC background, which has 71 % sequence similarity to SdcA. This finding suggests distinct physiological roles of SdcA and SidC, and a putative relationship between LotC and SdcA on regulating translation processes. Since LotC processes different types of ubiquitin chains, and is mainly localized to the cytosol, a catalytically dead version of LotC can be used as a standard tool for the identification of specific substrates of other known *Legionella* ligases. It would be interesting to understand how all these ubiquitin machineries synergistically work together and alter the host-ubiquitination system at different time points of infection.

## MATERIAL AND METHODS

### Protein expression and purification

All proteins used in this study were expressed and purified as previously described (Bhogaraju et al., 2016; Qiu et al., 2016). Lpg1621 (LotB), Lpg2529 (LotC), Lpg2411 and Lpg2907 were cloned into either pParallelGST2 or pParallelHis2 vector (Sheffield et al., 1999). T7 express *E. coli* competent cells (NEB) were transformed with plasmids and grown in LB medium to an OD_600_ of 0.6-0.8 at 37°C. Protein expression was induced by addition of 0.5 mM IPTG (isopropyl _D-thiogalactopyranoside_) and the cells were further grown overnight at 18°C and harvested. The cell pellet was resuspended in lysis buffer (50 mM Tris-HCl pH 7.5 and 150 mM NaCl, 2 mM DTT) and lysed by sonication and centrifuged at 13,000 rpm to clarify the supernatant. The supernatant of GST-tagged protein was incubated 1 hour with glutathione-*S*-sepharose pre-equilibrated with washing buffer (50 mM Tris-HCl pH 7.5, 500 mM NaCl, 2 mM DTT) and non-specific proteins were cleared with washing. GST-proteins were eluted with elution buffer (50 mM Tris-HCl pH 8.0, 50 mM NaCl, 2 mM DTT, 15 mM reduced glutathione) and buffer exchanged to storage buffer (50 mM Tris-HCl pH 7.5, 150 mM NaCl, 1 mM DTT). For His-tagged proteins, the supernatant was incubated with Ni-NTA pre-equilibrated with washing buffer (50 mM Tris-HCl, pH 7.5, 500 mM NaCl, 20 mM Imidazole) for 2 hours and eluted with elution buffer (50 mM Tris-HCl, pH 7.5, 500 mM NaCl, 300 mM Imidazole) and the buffer was exchanged to storage buffer. For LotC_14-310_, instead of using the elution buffer, glutathione beads were incubated with sfGFP-TEV protease (Wu et al., 2009) overnight at 4°C. Cleaved protein were buffer exchanged to IEX buffer A (20 mM Tris-HCl, pH 8.0, 20 mM NaCl, 1 mM DTT) and purified by anion-exchange chromatography on HitrapQ (GE Healthcare) with gradient elution with IEX buffer B (20 mM Tris-HCl, pH 8.0, 1 M NaCl, 1 mM DTT) and fractions contacting samples were loaded onto size-exclusion column (Superdex 75 16/60, GE Healthcare) pre-equilibrated with 50 mM Tris-HCl pH 7.5, 50 mM NaCl, 1 mM TCEP. Proteins were concentrated to 20 mg/ml and stored for crystallization.

### Di-Ub panel cleavage assay

To activate DUBs, 3 μl of 5 μM of DUBs were mixed with 12 μl of activation buffer (25 mM Tris-HCl pH 7.5, 150 mM NaCl, 10 mM DTT) and incubated 15 minutes at 25°C. For di-Ubiquitin samples, 3 μl of di-ubiquitin chains (1 mg/ml) were mixed with 3 μl 10x reaction buffer (500 mM Tris-HCl pH 7.5, 500 mM NaCl, 50 mM DTT) and 12 μl of ultra-pure water. To initiate the reaction, the activated DUBs were mixed with di-Ubiquitin and samples were taken at the indicated time points and the reactions were quenched by the addition of SDS-sample buffer. The samples were further analyzed by SDS-PAGE and stained with silver-staining kit (Pierce Silver Staining Kit, Thermo Fischer).

### Ubiquitin/NEDD8/SUMO-/ISG15/UFM activity-based probes assay

DUBs were diluted (1.5 μM, final concentration) with activation buffer and incubated 10 minutes at 25°C and the activity-based probes were diluted (50 μM, final concentration) in dilution buffer (50 mM Tris-HCl 7.5, 150 mM NaCl). The total of 30 ul of reaction mixture was prepared by mixing 20 μl of activated DUBs (1.5 μM), 3 μl of activity-based probes, 3 μl of reaction buffer (500 mM Tris-HCl pH 7.5, 500 mM NaCl, 50 mM DTT) and 4 μl of ultra-pure water. Samples were taken at the indicated time points and the reactions were quenched by the addition of SDS-sample buffer. Samples were further analyzed by SDS-PAGE and stained with Coomassie staining solution.

### Crystallization

The concentrated LotC_14-310_ were screened with sitting drop matrix screens in 96-well plate with 100 nl of protein and 100 nl of precipitant solution at 293K. Initial crystals appeared from solution containing 25 % PEG3350, 100 mM Tris-HCl pH 8.5, 200 mM NaCl with 18.4 mg/ml protein concentration. Diffraction-quality crystals were grown in optimized solution containing 19 % PEG 3350, 100 mM Tris-HCl pH 8.5, 150 mM NaCl with 24 mg/ml protein concentration.

### Data collection, processing and structure determination

To obtain the phase, 0.4 μl of 10 mM K_2_PtCl_4_ was added to the drop containing crystals and incubated for 18 hours. Heavy atom-soaked crystals were cryo-protected using mother liquor solution supplemented with 15 % (v/v) glycerol. Diffraction data were collected on single frozen crystal in a nitrogen stream at 100K at beamline PXI as Swiss Light Source, Villigen. Initial data sets were processed using XDS (Kabsch, 2010), and initial-phases were determined by Autosol in Phenix (Terwilliger et al., 2009). Structure refinement and manual model building were performed with Coot and Phenix.Refine (Afonine et al., 2012; Emsley et al., 2010).

### Protein-protein docking

We used the Rosetta protein-protein docking method (Gray et al., 2003) to identify low-energy conformations of the complexes of ubiquitin with LotB and LotC, respectively. Given that the C-terminus of ubiquitin should interact with the catalytic residue of the OTUs, we used the local docking approach in which we placed the C-terminal end of ubiquitin (Gly76) near the catalytic residues in both ligases (Cys29 and His270 for LotB and Cys16 and His296 for LotC). We then started the docking by optimizing the rigid-body orientation and side-chain conformation sampling. The program requires two protein structures as inputs, which were prepared by running the refinement protocol before the docking step. We performed the local docking approach and generated 100 independent structures for each complex. The complexes in this way were subject to local refinement to remove remaining small clashes. The complexes were then clustered based on the distance matrix of Cα atoms between the ligase and ubiquitin using the KMeans method. The representatives of two major clusters in each case were selected based on the interface score (I_sc), which represents the energy of the interactions across the interface of two proteins. These representative complexes were used for MD simulations.

### Molecular dynamics simulations

All-atom explicit solvent molecular dynamics (MD) simulations were performed for two docking results for each ligase. The systems were built using the CHARMM-GUI webserver (Wu et al., 2014). The systems were hydrated with 150 mM NaCl electrolyte. The all-atom CHARMM36m force field was used for proteins, lipids and ions, and TIP3P was used for water molecules (Best et al., 2012). The MD trajectories were analyzed with Visual Molecular Dynamics (VMD) (Humphrey et al., 1996). The MD simulations were performed using GROMACS 2019 (Abraham et al., 2015). The starting system was minimized for 5,000 steps with the steepest descent energy minimization and equilibrated for 6.5 ns of MD simulation first in the NVT ensemble (1.5 ns) and then in the NPT (5 ns) ensemble, in which all non-hydrogen atoms of the protein were restrained to the fixed reference positions with progressively reduced force constants, starting at 1000 kJ·mol^−1^·nm^2^. Afterwards, the production runs were carried out in the NPT ensemble for 600 ns for each setup. To keep the C-terminusof ubiquitin in the catalytic site, 7-Å wall restraints were placed on the distance between Cα of G76_UB_ and Cys29/His270 in LotB and between Cα of G76_UB_ and Cys16/His296 in LotC. Periodic boundary conditions were used. Particle mesh Ewald (Darden et al., 1993) with cubic interpolation and 0.12 nm grid spacing for Fast Fourier Transform was used to treat long-range electrostatic interactions. The time step was 2 fs. The LINCS algorithm (Hess et al., 1997) was used to fix all bond lengths. Constant temperature (310 K) was set with a Nosé-Hoover thermostat (Hoover, 1985), with a coupling constant of 1.0 ps. An isotropic Parrinello-Rahman barostat (Parrinello and Rahman, 1981) with a coupling constant of 5.0 ps was used to maintain a pressure of 1 bar.

### Identification of novel substrates of LotB and LotC

For interactome analysis, HEK 293 cells were transfected with FLAG-LotB WT/C29A or FLAG-LotC WT/C24A. To identify the substrates modified by *Legionella* derived E3 ligases, GFP-SidC or GFP-SdcA were co-transfected with FLAG-LotB WT/C29A or FLAF-LotC WT/C24A. Cells were lysed in ice cold lysis buffer (50mM Tris-Cl, p.H 7.5; 150mM NaCl; 1% Triton x-100) and equal amount of lysates were incubated with FLAG-M2 beads in IP buffer (Lysis buffer without detergent). After incubation, IPs were washed with wash buffer (50mM Tris-Cl, pH7.5; 400mM NaCl; 0.5mM EDTA) and the interacting proteins were eluted with 8M urea solution. After the reduction and alkylation with TCEP and Chloroacetamide, the samples were digested with 0.5ug Trypsin (Promega) at 37°C overnight after diluting the urea < 2M. Digests were acidified using trifluoroaceticacid (TFA) to a pH of 2-3, and the peptides were enriched using C18 stage tips (Rappsilber et al., 2003). To get quantitative information, peptides were either labelled with TMT 10 plex reagent (Thermo fisher) or analyzed label-free. The peptides were separated on an in-house made C18 column (20 cm length, 75 μm inner diameter, 1.9 μm particle size) by an easy n-LC 1200 (ThermoFisher) and directly injected in a QExactive-HF or in case of TMT samples into a Fusion Lumos mass-spectrometer (ThermoFisher) and analyzed in data-dependent mode. Data analysis was done using Maxquant 1.65 (Cox and Mann, 2008). Fragment spectra were searched against *Homo sapiens* SwissProt database (TaxID:9606, version 2018). Label-free quantification was done with MaxLFQ (Cox et al., 2014) method with activated match between runs. TMT-labelled samples were analysed by using TMT 10 Plex option within the software. Statistically significant changes between samples were determined using a Two-sample t-test with a permutation-based FDR of 5% on log2 transformed values in Perseus (Tyanova et al., 2016).

### Confocal imaging and image analysis

U2OS cells were transfected with FLAG-LotB or FLAG-LotC by GeneJuice transfection reagent (Merck) for 24 h and fixed by 4% paraformaldehyde for 20 min. After fixation, cells were permeabilized and blocked by 0.1% saponin and 1% BSA in PBS for 1 h at room temperature. Cells were incubated with anti-Flag antibody (Sigma and Cell Signaling), with either anti-calnexin antibody (Abcam), anti-TOMM20 antibody (Abcam), or anti-GM130 (BD Transduction Laboratories) at 4 ℃ overnight. Alexa Fluor 488 and Alexa Fluor 546 (Invitrogen) secondary antibody were incubated for 1 h at room temperature. Images were acquired by the Zeiss LSM780 microscope system with 63× 1.4 NA oil immersion objective and further analyzed by Zeiss Zen microscope software.

## ACKNOWLEDGEMENTS

We thank Yuxin-Mao for providing SidC and SdcA clones and Stefan Knapp for the advice in structure determination and sharing synchrotron time. The authors also thank staff at SLS for their support during crystallographic X-ray diffraction test and data collection. The data collection at SLS has been supported by the funding from the European Union’s Horizon 2020 research and innovation program under grant agreement number 730872, project CALIPSOplus. This project was supported by the European Research Council (ERC) under the European Union’s Horizon 2020 research and innovation program (ID, grant agreement No 742720), the LOEWE program DynaMem of the State of Hesse (Germany, Project-ID III L6-519/03/03.001 – (0006)), and Deutsche Forschungsgemeinschaft (DFG, German Research Foundation (Project-ID 259130777 – SFB1177; Leibniz-Program to ID; CEF-MC - EXC115/2; SFB 902), the Max Planck Society and NWO-VIDI grant and Off-rad grand for G.H.

## COMPETING INTERSTES

The authors declare no competing interests.

---

**Supplementary Figure 1.**
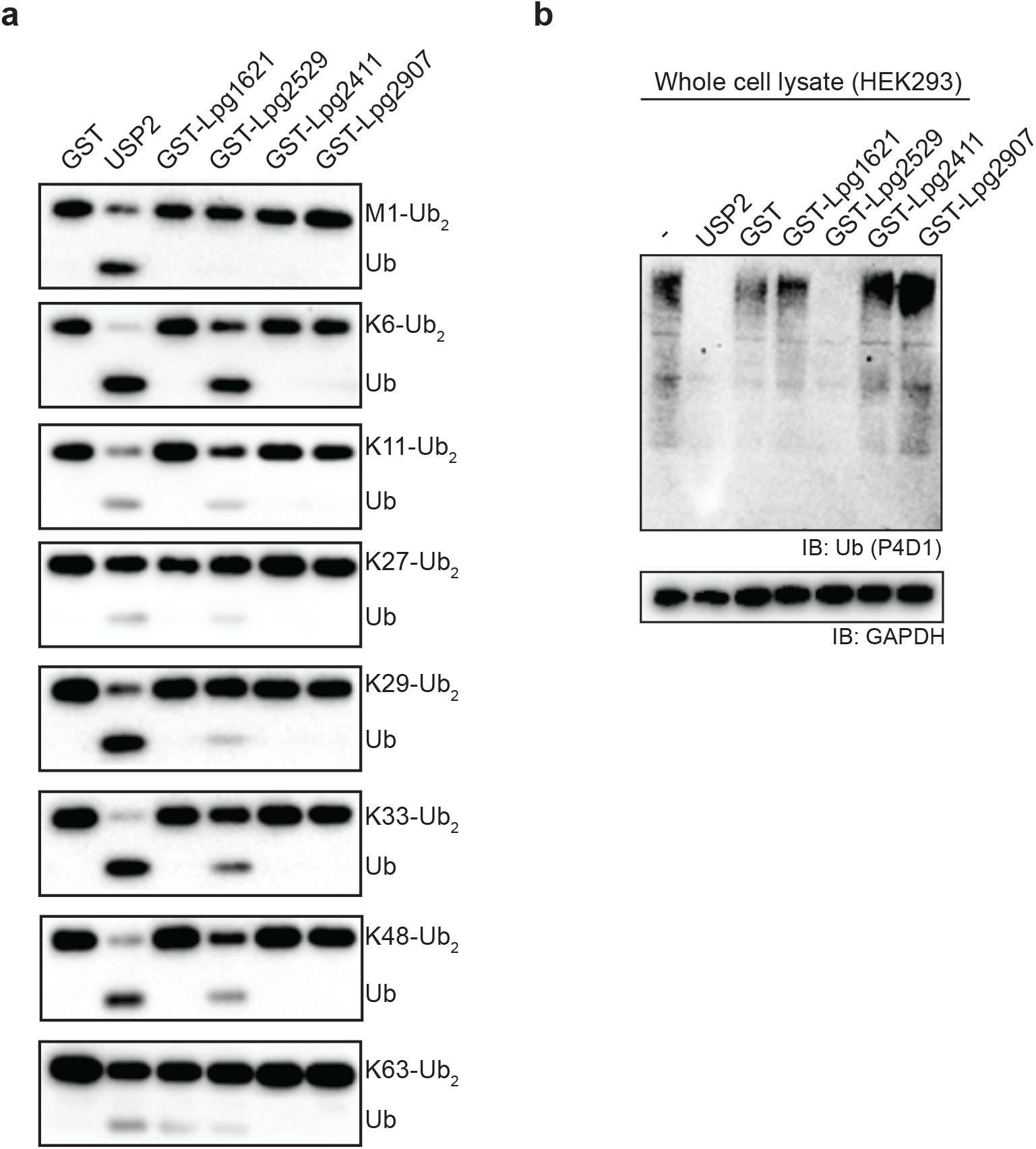

**Supplementary Figure 2.**
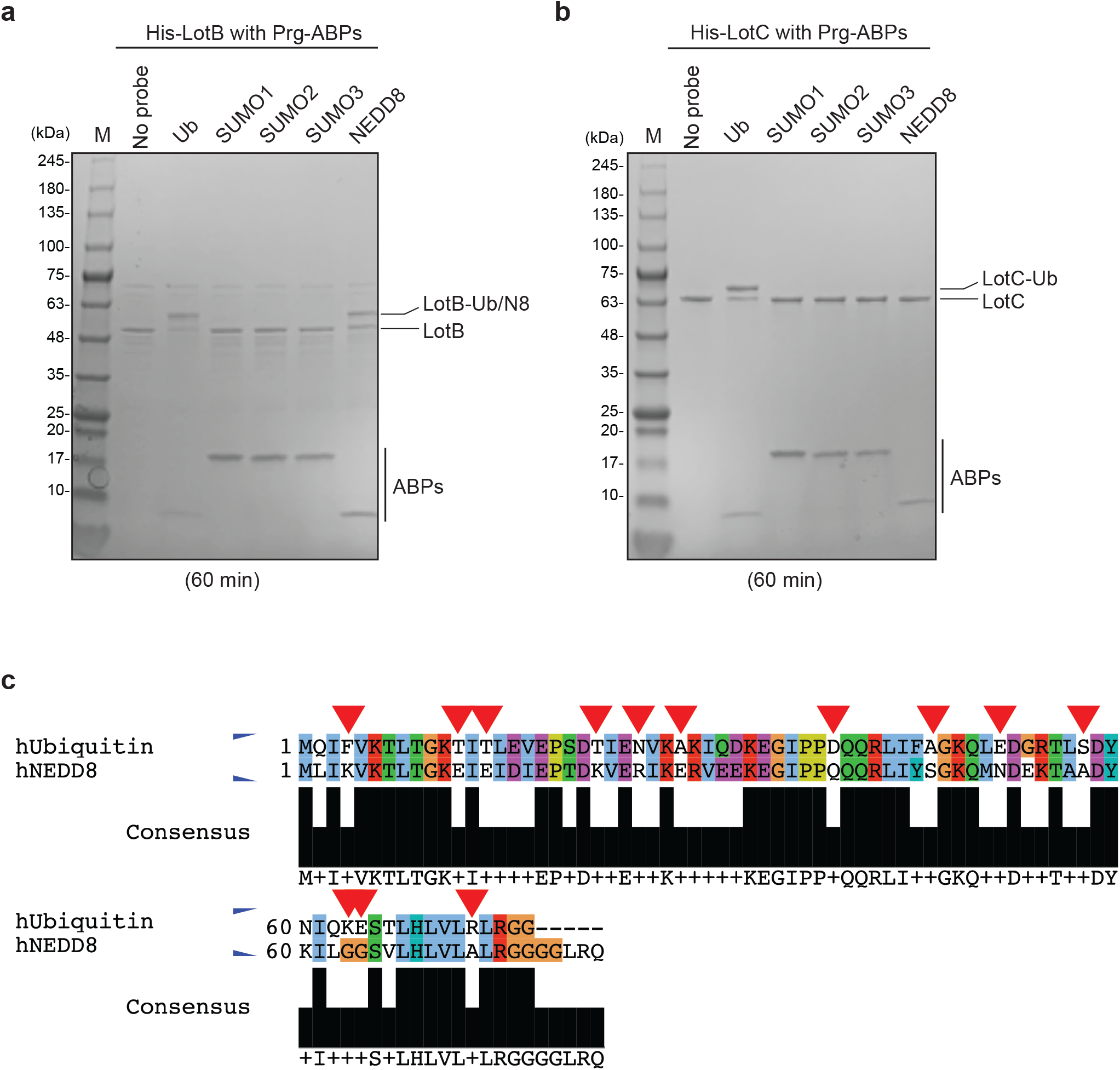

**Supplementary Figure 3.**
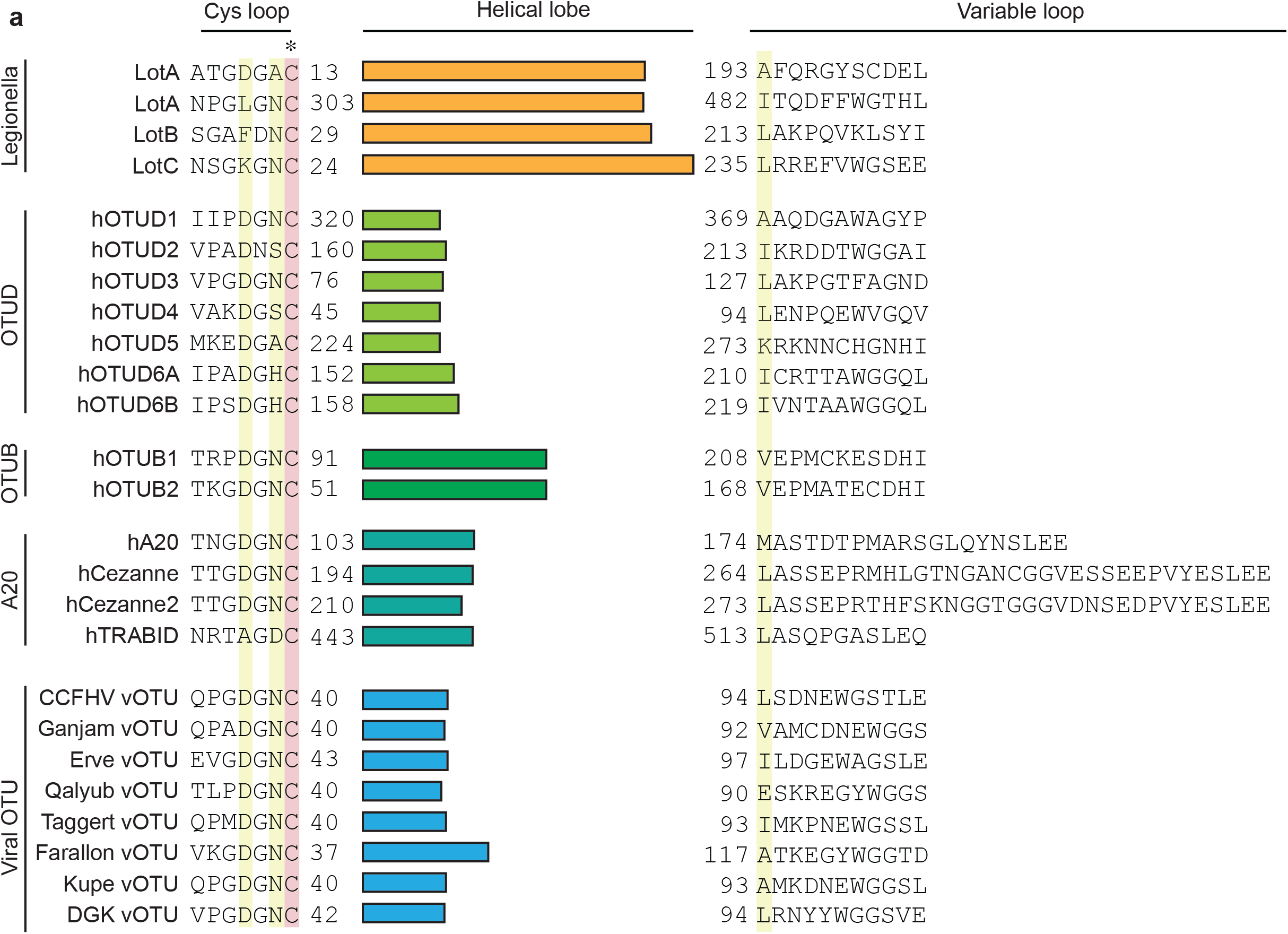

**Supplementary Figure 4.**
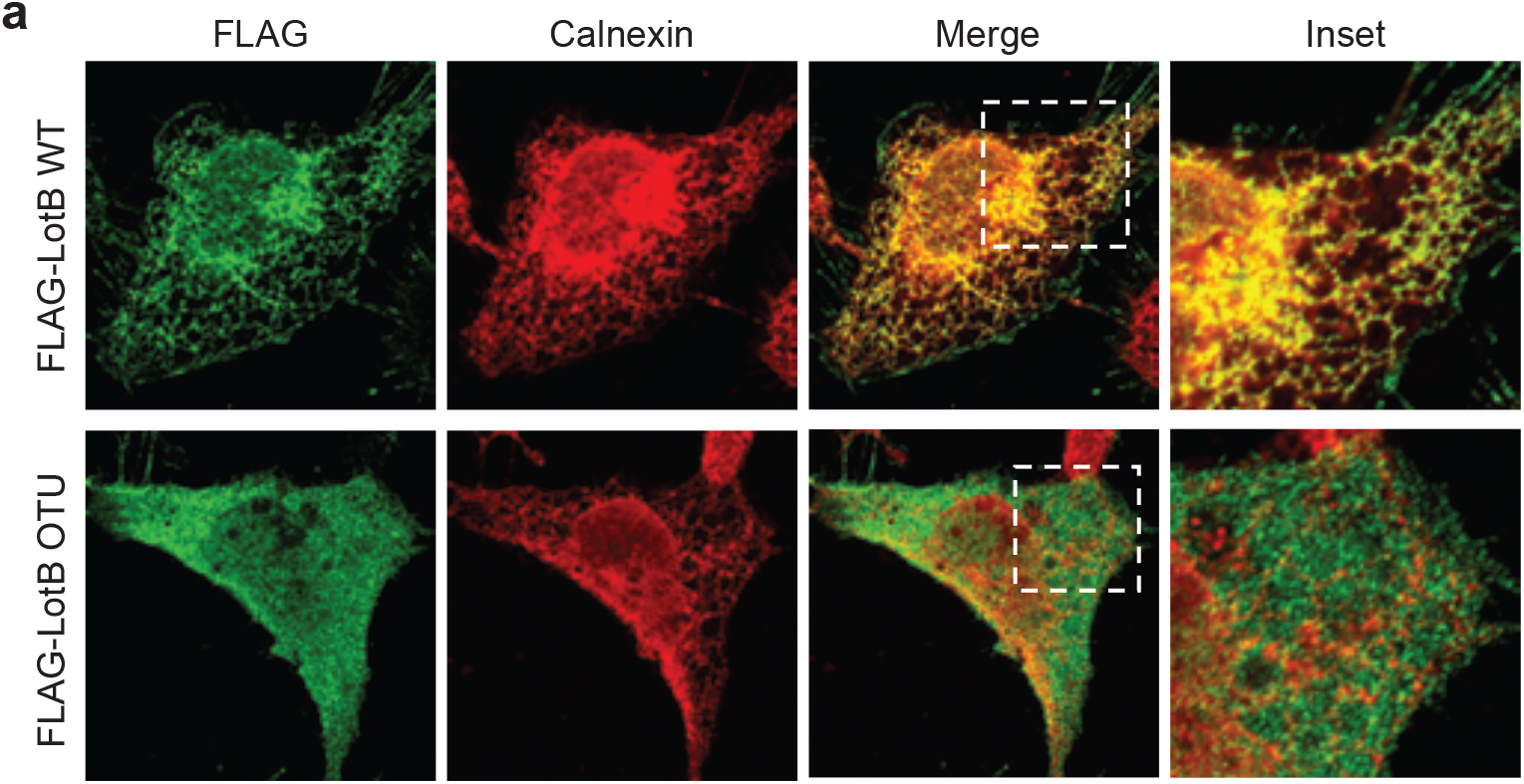

**Supplementary Figure 5.**
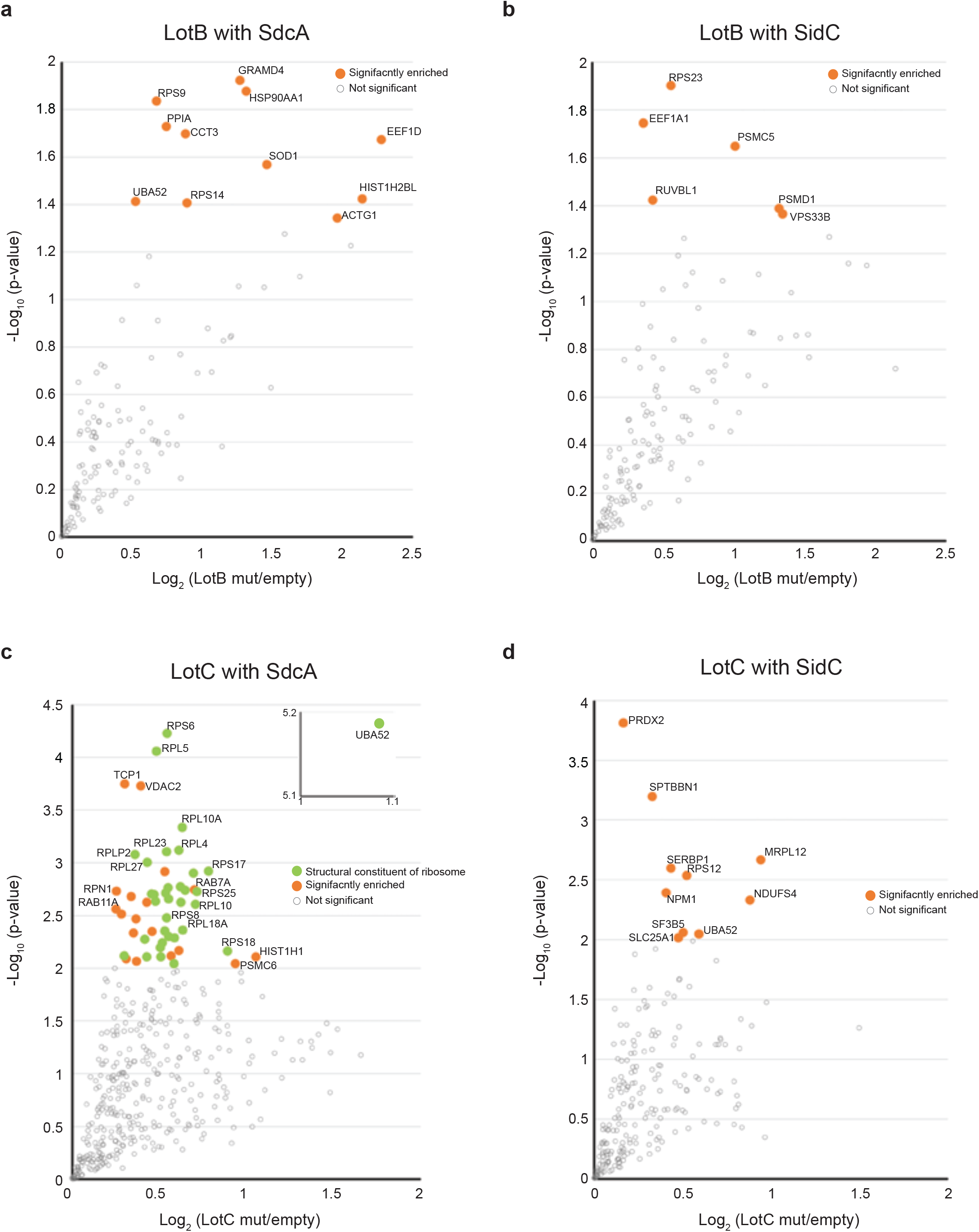

**Supplementary Figure 6.**
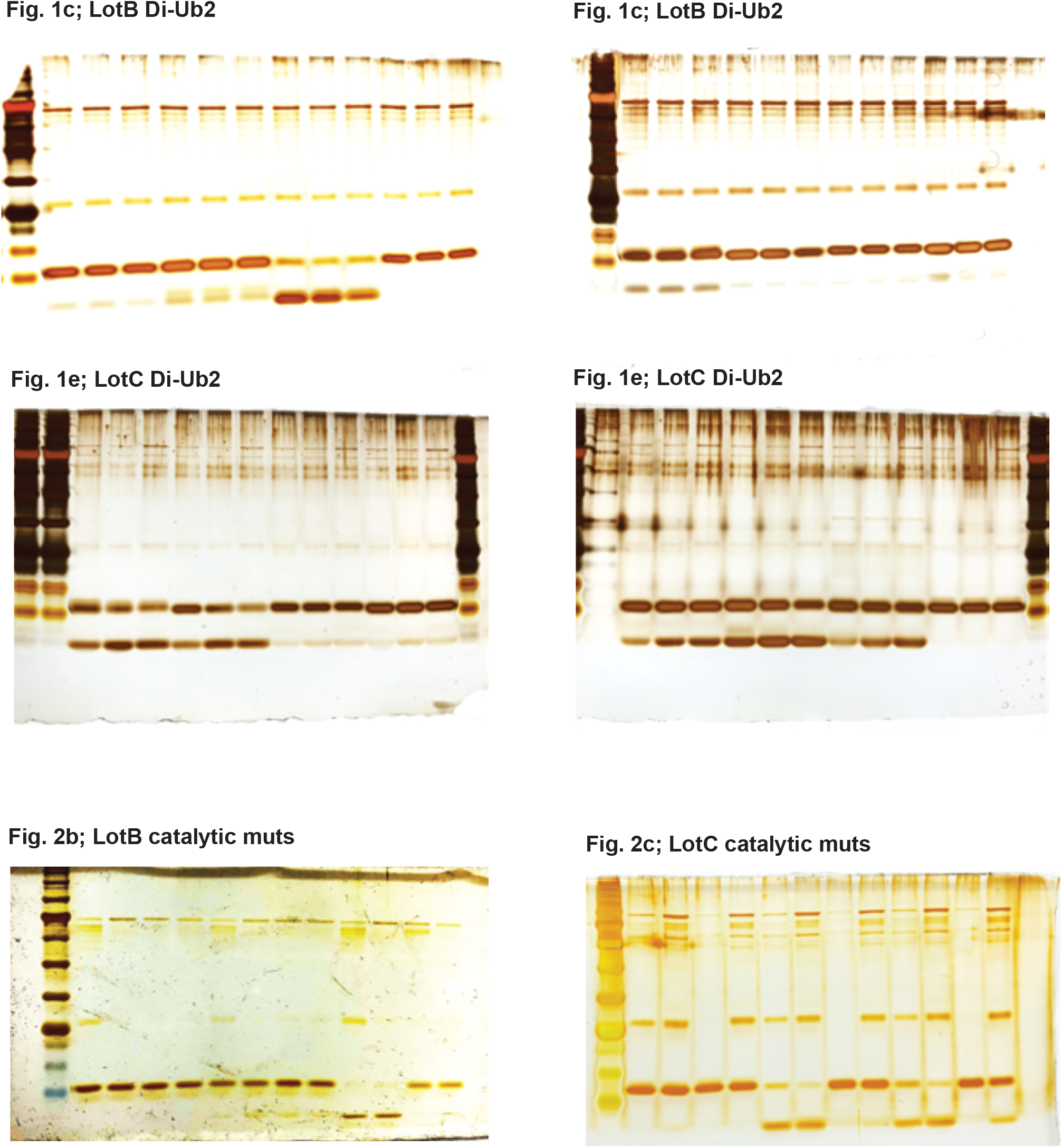
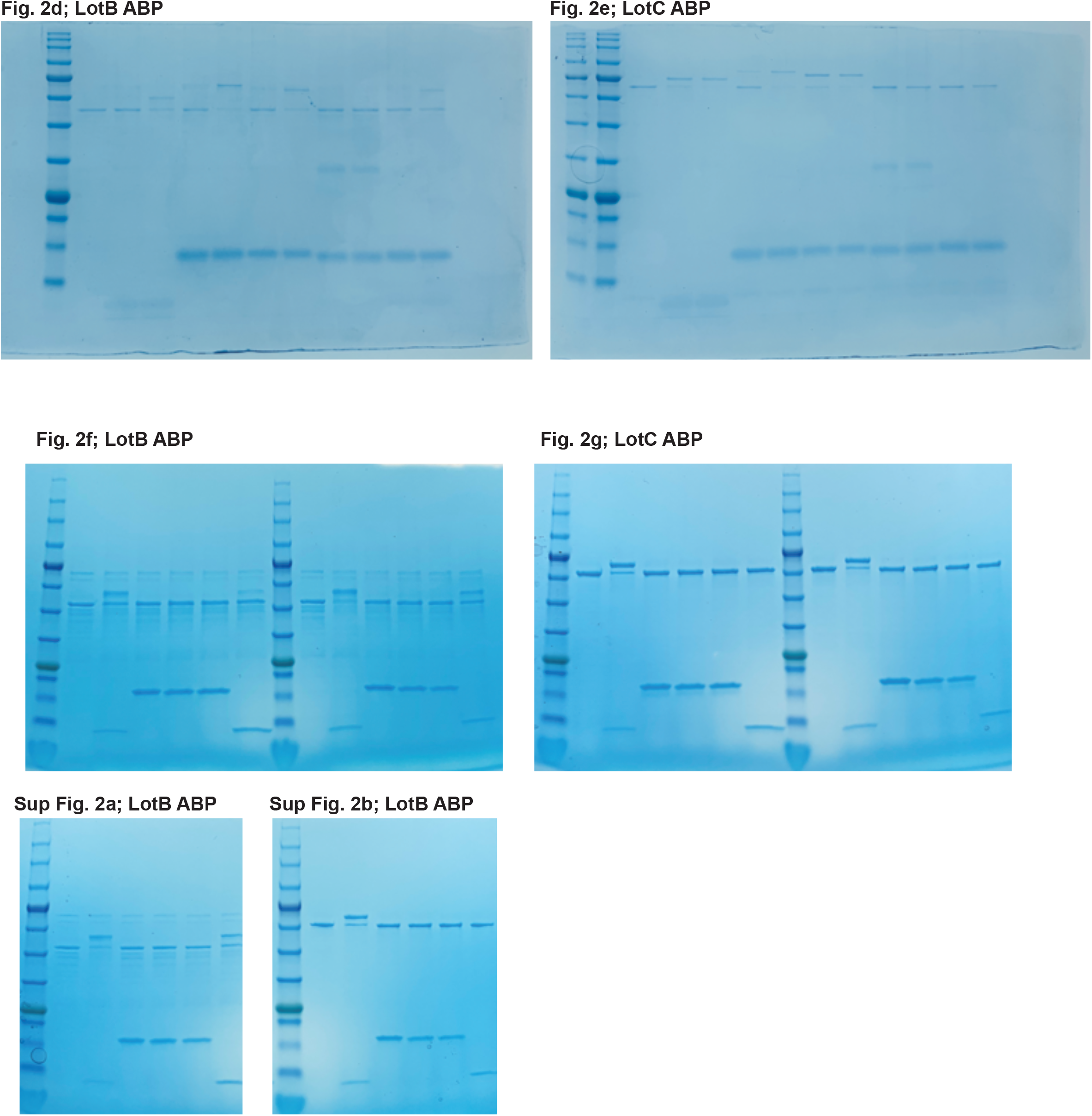
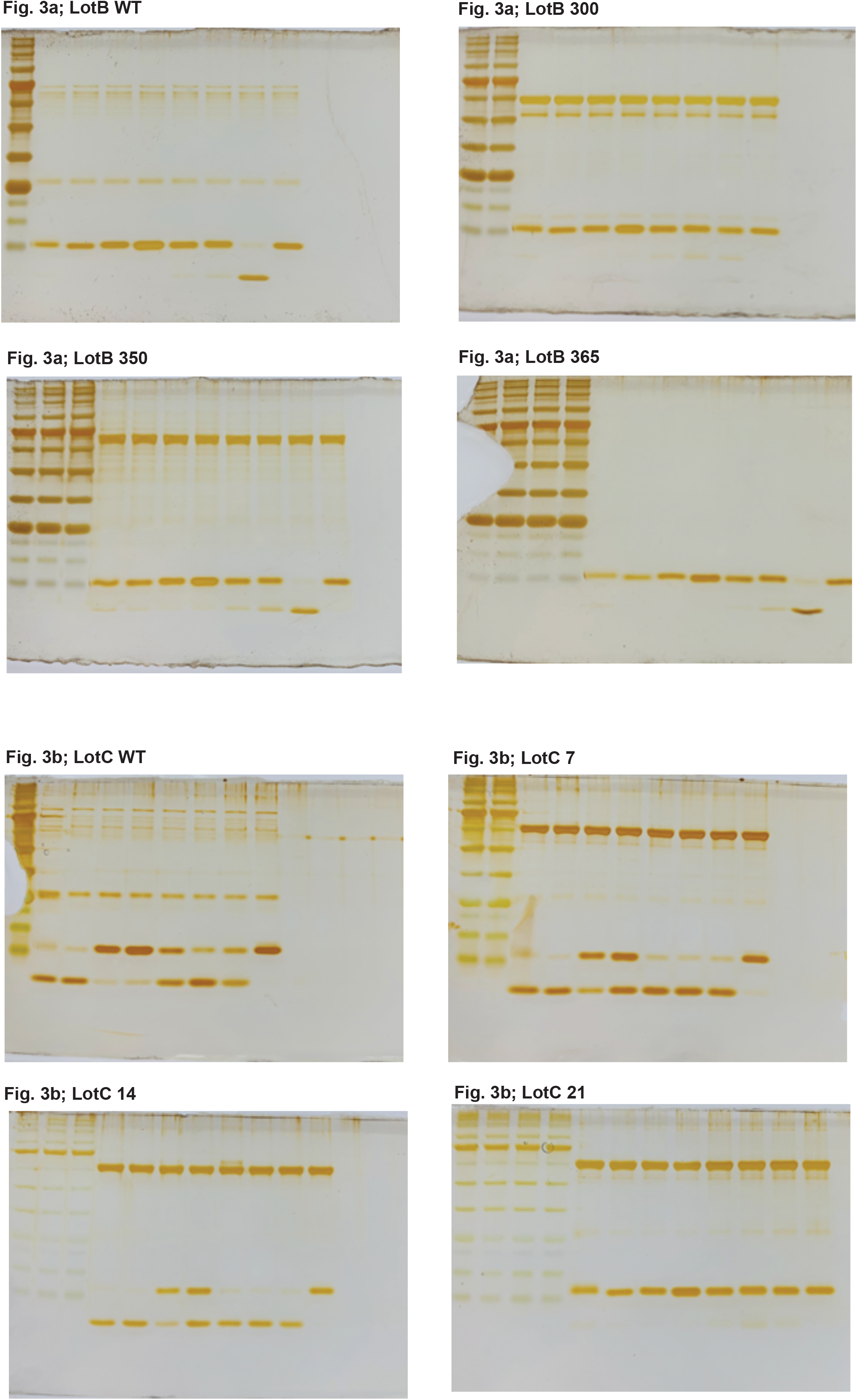
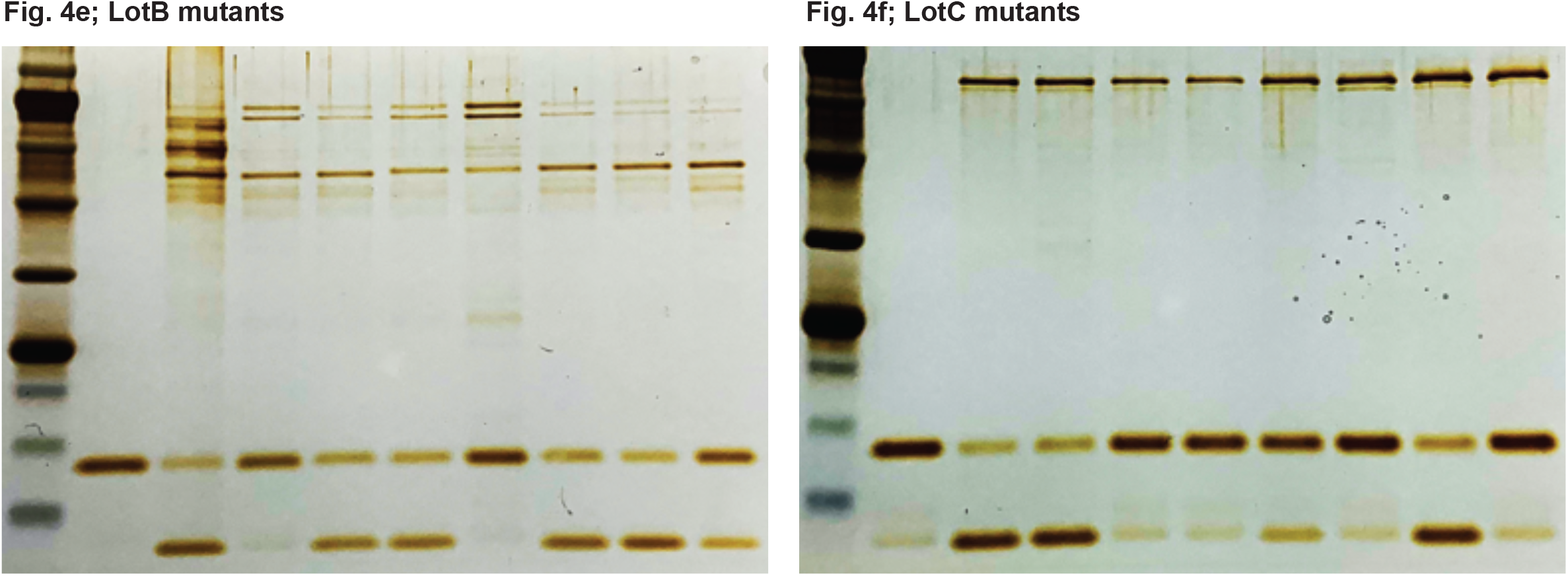
Original blots

